# Nitrative signaling into cardiac lactate dehydrogenase Trp^324^ modulates active site loop mobility and activity under metabolic stress

**DOI:** 10.1101/348789

**Authors:** Cristian Nogales, Jeroen Frijhoff, Detlef Thoenges, Richard B. Sessions, Albert Sickmann, Tatiana Nedvetskaya, Cesar Ibarra-Alvarado, Vera Schmitz, Peter Rösen, Axel Goedecke, Aleksandar Jovanović, David H. Perlman, Anna Klinke, Stephan Baldus, Ana I. Casas, Péter Bai, Csaba Szabo, Martin Feelisch, Harald H.H.W. Schmidt

**Affiliations:** Department of Pharmacology and Personalised Medicine, Maastricht University, Maastricht, The Netherlands; Medtronic Bakken Research Center, Maastricht, The Netherlands; DiaSys Diagnostic Systems GmbH, Holzheim, Germany; School of Biochemistry, University of Bristol, Bristol, UK; Leibniz-Institut für Analytische Wissenschaften – ISAS, Dortmund, Germany; Rudolf-Buchheim-Institue for Pharmacology, Justus-Liebig-University, Giessen, Germany; School of Chemistry, Autonomous University of Querétaro. Querétaro, México; German Diabetes Research Institute, Düsseldorf, Germany; Heinrich-Heine-University, Department Cardiovascular Physiology, Düsseldorf, Germany; University of Nicosia Medical School, Nicosia, Cyprus; Merck Exploratory Science Center, Cambridge, Massachusetts; Cologne Cardiovascular Research Center CCRC, Cologne, Germany; Department of Medical Chemistry, Faculty of Medicine, University of Debrecen, Debrecen, Hungary; MTA-DE Lendület Laboratory of Cellular Metabolism, Debrecen, Hungary; Department of Oncology, Microbiology and Immunology; University of Fribourg, Fribourg, Switzerland; University of Southampton Medical School, Southampton, UK

**Author notes:** Both first authors contributed equally to this work (Cristian Nogales, Jeroen Frijhoff). **One Sentence Summary**: We here expand protein nitration from mainly affecting tyrosine moieties and being a disease marker of oxidative stress to physiological nitration of a tryptophan in cardiac lactate dehydrogenase downregulating its maximal activity under basal conditions and metabolic stress.

## Abstract

Protein tyrosine nitration is a hallmark of oxidative stress related disease states, commonly detected as anti-nitrotyrosine immunoreactivity. The precise reactive oxygen sources, mechanisms of nitration as well as the modified target proteins and functional consequences, however, remain often unclear. Here we explore protein tyrosine nitration under basal conditions and find surprisingly physiologically nitrated proteins. Upon purifying a prominent physiologically nitrotyrosine immunopositive in hearts from mouse, rat and pig, we identify it as lactate dehydrogenase (LDH). Mechanistically, LDH’s degree of basal nitration depended on two canonical sources, NO synthase (NOS) and myeloperoxidase (MPO), respectively. When validating the nitrated amino acid by MALDI-TOF mass spectrometry, we, surprisingly, located LDH nitration not to a tyrosine but the C-terminal tryptophan, Trp^324^. Molecular dynamics simulations suggested that Trp^324^ nitration restricts the interaction of the active site loop with the C-terminal α-helix essential for activity. This prediction was confirmed by enzyme kinetics revealing an apparent lower V_max_ of nitrated LDH, although yet unidentified concurrent oxidative modifications may contribute. Protein nitration is, thus, not a *by definition* disease marker but reflects also physiological signaling by eNOS/NO, MPO/nitrite and possibly other pathways. The commonly used assay of anti-nitrotyrosine immunoreactivity is apparently cross-reactive to nitrotryptophan requiring a reevaluation of the protein nitration literature. In the case of LDH, nitration of Trp^324^ is aggravated under cardiac metabolic stress conditions and functionally limits maximal enzyme activity. Trp^324^-nitrated LDH may serve both as a previously not recognized disease biomarker and possibly mechanistic lead to understand the metabolic changes under these conditions.

## Introduction

Increased levels of reactive oxygen species (ROS), i.e. oxidative stress, is considered a common mechanism of several cardiovascular and other disease states (*1*). In concert with different nitrogen species (*2*–*4*), ROS can lead to protein nitration, e.g. tyrosine residues (NO_2_Tyr). Tyrosine nitration is the most frequently used and considered the most robust disease marker of oxidative stress (*5*–*7*). Its detection relies mainly on NO_2_Tyr specific antibodies (*8*–*10*). With respect to the nitrogen source, anti-NO_2_Tyr immunopositive signals are generally interpreted as a hallmark of nitric oxide (NO) being scavenged by superoxide and intermediate peroxynitrite (*9*, *11*, *12*), although other mechanisms exist (*13*). Regarding the functional consequences of protein nitration few *in vitro* data exist suggesting mainly loss of protein function correlating with disease severity (*14*–*17*). *In vivo* data are lacking. Moreover, little is known about the protein factors that govern other nitration sites, e.g. tryptophan, and about the effects of the modification in protein structure and function (*18*). Here, we address three major knowledge gaps in our understanding of protein nitration: specificity, origin, and disease relevance.

## Results

### Development of an anti-nitrotyrosine antibody biomarker panel for disease-relevant oxidative stress

We started out by examining a panel of anti-NO_2_Tyr antibodies (Fig. 1), including those directed against carrier-linked NO_2_Tyr or against H_2_O_2_/NO_2_^−^ treated proteins (Table S1). We tested these in rodent (rat) and, for later purification, also in porcine tissues. Surprisingly we obtained signals already under physiological conditions. The immunoreaction, however, varied greatly in protein band size and intensity. Two polyclonal antibodies (pAb), pAb1 and pAb2, and one monoclonal antibody (mAb), mAb1, displayed the most robust signals (Fig. S1). The specificity of anti-NO_2_Tyr antibodies (*19*–*23*) is rarely validated, e.g. by antibody pre-absorption with free NO_2_Tyr (*11*). In our case, pAb1 signals were partially blocked by pre-absorption with 3 mM NO_2_Tyr. mAb1 showed very little non-specific background whilst pAb2 exhibited moderate sensitivity with high anti-NO_2_Tyr specificity.

**Fig. 1.**
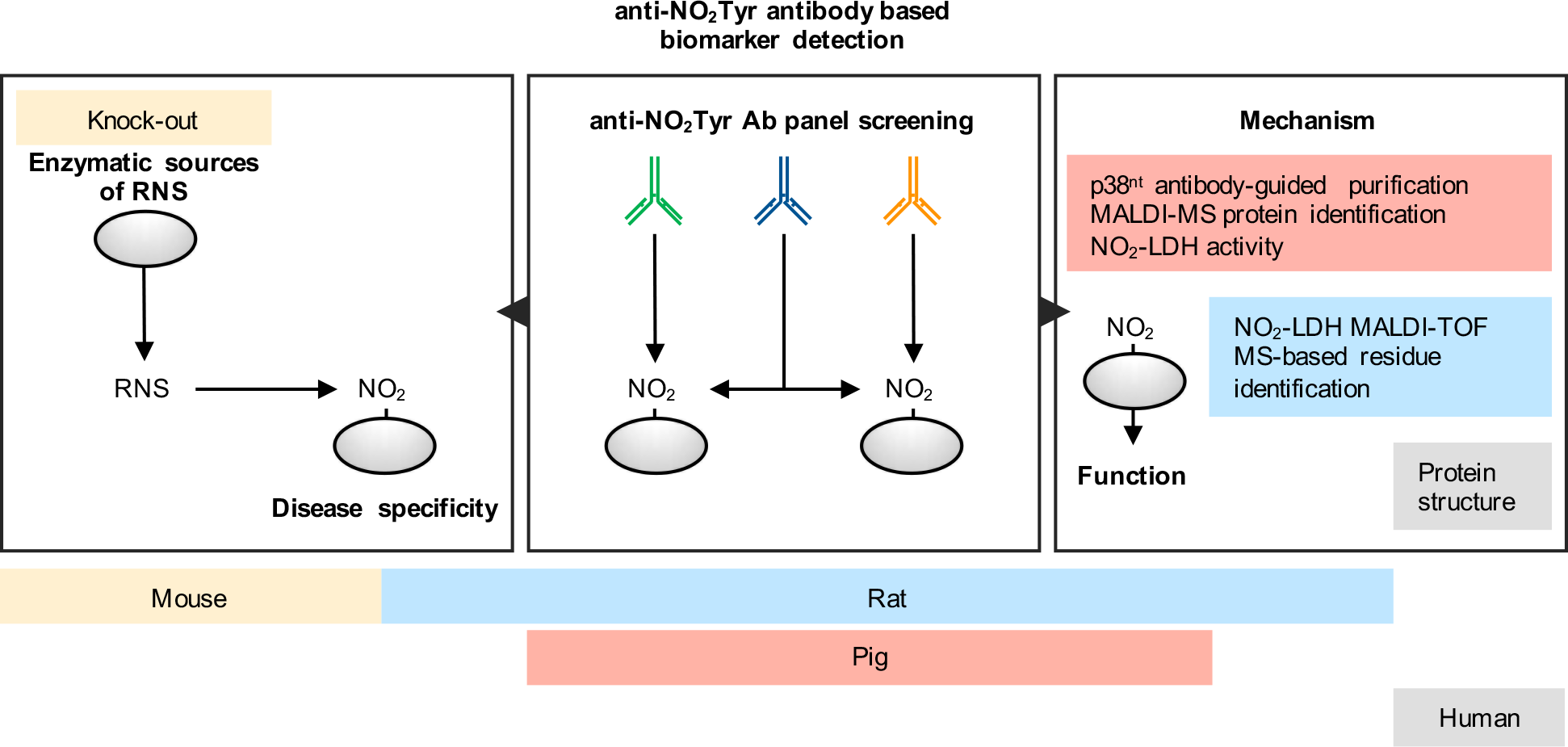
Experimental outline for an anti-NO_2_Tyr antibody-based biomarker detection approach. Central panel: Antibody panel screening, an anti-NO_2_Tyr antibody panel was tested in porcine and rat tissues under physiological conditions. Different antibodies might recognize different nitrated proteins (green and orange antibodies) or one antibody might detect more than one nitrated protein (blue antibody). Right panel: A 38 kDa band was especially prominent and, subsequently, purified in large mammals and identified via MALDI-TOF as LDH. NO_2_-LDH activity was also assessed. We identified the nitrated residue and simulated the effect of the nitration in the protein structure and mechanism using human LDH. Left panel: Different rodent models, as well as knock-out mouse models were used to elucidate the effects of NO_2_-LDH in disease, diabetes and myocardial stress, and the enzymatic sources of reactive nitrogen species (RNS) leading to LDH nitration, respectively.

### A prominent nitrotyrosine immunoreactive protein band was detected in mouse, rat and pig heart under physiological conditions

To conduct a tissue/species screen, we selected three antibodies, pAb1, pAb2, and mAb1, from our original panel. Three proteins were consistently detected under basal conditions and with apparent molecular weights of 70, 45 and 38 kDa, of which the latter gave the by far the strongest signal in heart (Fig. 2A). Pre-incubation with free NO_2_Tyr or NO_2_BSA (3 mM) or nitro-group reduction (*22*), at least partially, blocked the pAb1 dependent signal. Therefore, subsequent efforts were directed at identifying the nature of the NO_2_Tyr positive 38 kDa protein and its site of nitration.

**Fig. 2.**
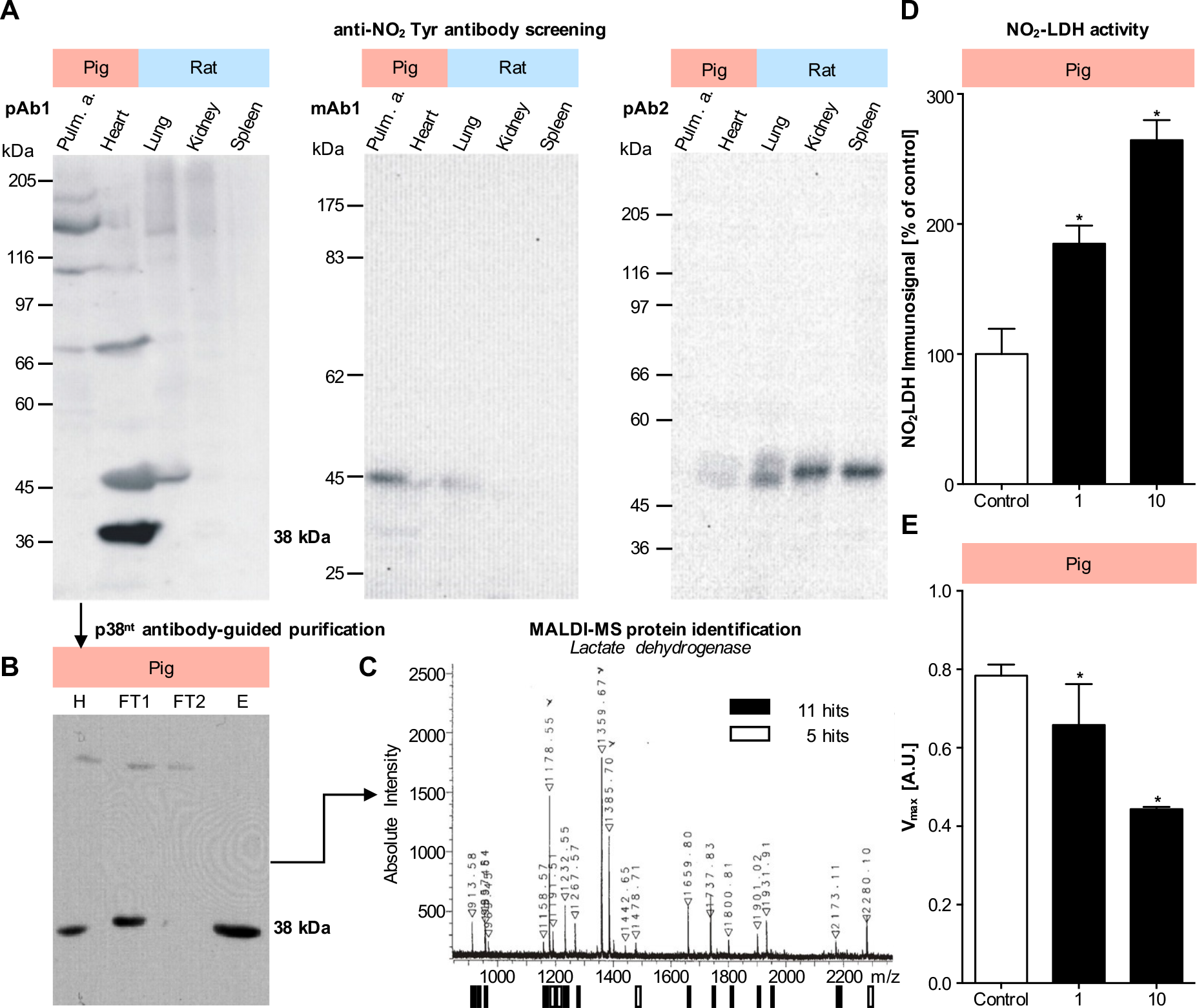
Screening of different native tissue homogenates identifies a 38 kDa band in heart as major physiologically NO_2_Tyr-immuno positive protein. Further purification and MALDI-MS analysis identified the 38 kDa band as LDH. (**A**) Immunopositive pig and rat tissues from Fig S1 were further analyzed by SDS-PAGE under reducing conditions (dithiothreitol) and subsequent western blot. The band pattern was different for all three polyclonal (pAb) or monoclonal (mAb) antibodies: pAb1, mAb1 and pAb2. The most intense NO_2_Tyr immunopositive protein band at approximately 38 kDa (p38^nt^) was detected using pAb1 in heart. (**B**) p38^nt^ was purified from porcine heart homogenate by affinity (Blue Sepharose) followed by ion-exchange (Q-Sepharose) chromatography. Purification steps were tracked via protein nitrotyrosine detection using pAb1. (**C**) Purified cardiac p38^nt^ was cut out from the gel, digested with trypsin and analyzed by mass spectrometry and peptide mass fingerprint analysis. Most peptide ions detected were matched to lactate dehydrogenase: 11 ions matched to LDH (∎) out of a total of 20 detected, with 5 ions matching to the next best protein assignment, malate dehydrogenase (MDH, ☐), identifying p38^nt^ as LDH with a p<0.05. (**D**) LDH nitration was assessed by quantitative western blot analysis using pAb1 (n=9). (**E**) Vmax[LDH] remained unchanged up to an equimolar peroxynitrite:LDH ratio, and then decreased at a ratio of 10 (n=8). Data represent means ±SEM.

### Upon purification, the 38 kDa nitrated protein was identified as LDH with lower V_max_

Using porcine heart, we isolated the NO_2_Tyr immunoreactive 38 kDa protein by ion exchange (Q Sepharose, QS) and affinity (Blue Sepharose, BS) chromatography (Fig. 2B). The partially purified 38 kDa protein was stained by colloidal Coomassie Blue and subjected to peptide mass fingerprinting using MALDI-TOF MS. Based on the NCBIprot database most ions present in the spectrum of tryptic peptides matched LDH as the primary protein component of the sample (Fig. 2C).

Next, we tested the effect of nitration on the enzyme activity by nitrating cardiac LDH *in vitro* with increasing amounts of H_2_O_2_/NO_2_^−^. Indeed, higher nitration levels of LDH (NO_2_-LDH) (Fig. 2E) correlated with decreased V_max_ (Fig. 2F) and unchanged K_M_. Our data suggest therefore that protein nitration is not only a physiological event for cardiac LDH, but also an *in vivo* mechanism modulating enzyme activity by lowering V_max_.

### Nitrated LDH is increased in diabetes and myocardial stress

To investigate whether NO_2_-LDH is further increased in response to metabolic stress, we investigated heart extracts from rat models of streptozotocin-induced short-term (ST) and long-term (LT) diabetes mellitus (*24*) as well as a doxorubicin (DoxoR) induced mouse model of cardiac stress (*25*) (Fig. 3A). Diabetic rats were sacrificed 3 (ST) and 16 (LT) weeks after inducing diabetes at week 10. Using the pAb1, NO_2_-LDH signals were significantly increased in all three models when compared against healthy controls (Fig. 3, B and C). Immunosignals of pAb1 pre-absorbed with NO_2_Tyr or anti-LDH wildtype antibody did not show any significant change. In both diabetes mellitus models and doxorubicin-treatment animals, levels of cardiac LDH were unchanged versus controls, indicating that indeed the NO_2_-LDH/LDH ratio was increased.

**Fig. 3.**
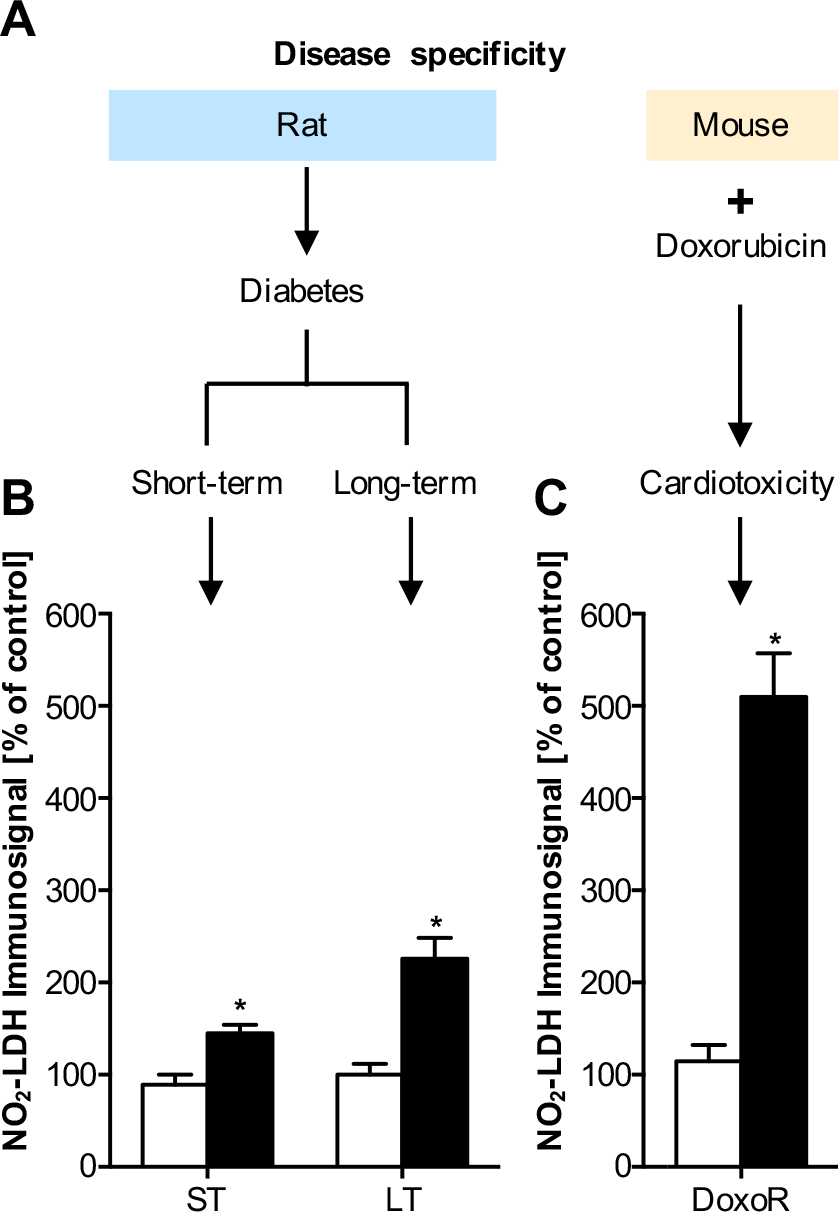
Pathological LDH nitration levels in short-term (ST) and long-term (LT) diabetic disease models and an oxidative stress myocardial dysfunction model. (**A**) Heart homogenates of ST and LT streptozotocin-treated diabetic rats or doxorubicin-treated mice (DoxoR) were analyzed by SDS-PAGE and immunoblot analysis for LDH nitration using the pAb1 antibody. In all three animal models, the relative NO_2_-LDH signal intensity was increased. In diabetes, the long-term model yielded higher values than the short-term model, which may indicate either higher nitrosative stress, or, more likely, accumulation of LDH nitration (**B**). The highest increase was observed in the doxorubicin model (**C**). Data are expressed as % of untreated control animals (open bars) and represent means ± SEM of n=6-18 experiments.

### Cardiac LDH nitration is lowered in NO synthase (eNOS) knockout (KO) and myeloperoxidase (MPO) KO mice

We next wanted to address our second goal, to understand the mechanistic origin of the nitrogen in NO_2_-LDH. Usually, it is assumed that protein nitration derives from peroxynitrite from the reaction between ^•^NO and O_2_^•−^ (*26*). However, another major source, not involving de novo NO synthesis is frequently overlooked, i.e. MPO-catalyzed nitration using nitrite in the presence of hydrogen peroxide (*27*) (Fig. 4A). To analyze this *in vivo*, we had to switch to mouse models, i.e. eNOS-KO and MPO-KO lines. Indeed, we observed that NO_2_-LDH signals in heart tissue were not only reduced in eNOS-KO but also in MPO-KO mice when compared to their respective WT litter mates (Fig. 4B). These results therefore indicated that both pathways contribute to basal nitration of cardiac LDH. Moreover, we also compared LDH nitration to oxidation and showed that oxidation occurred at lower peroxynitrite concentrations and was half-maximal when LDH activity was still unchanged (Fig. S2).

**Fig. 4.**
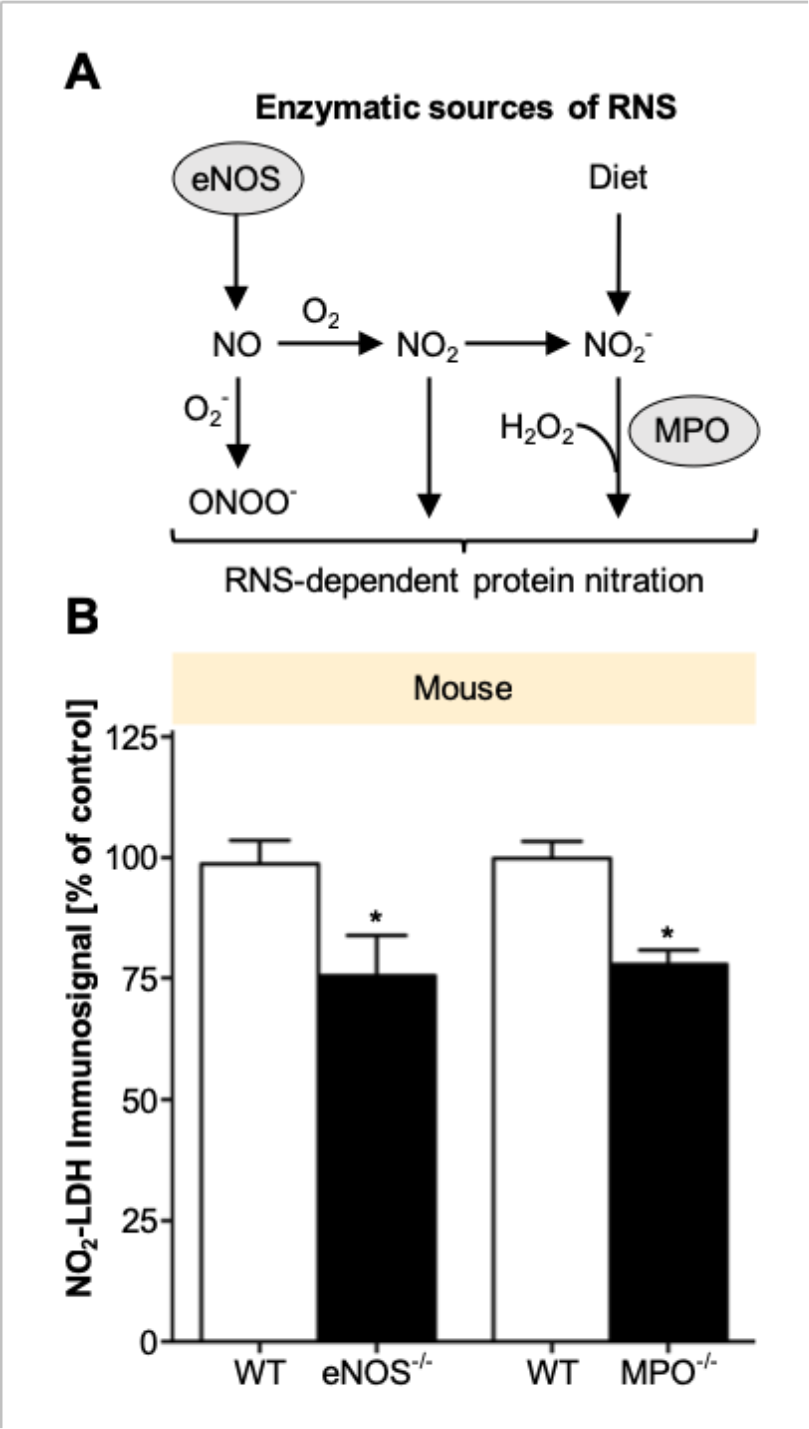
*In vivo* genetic validation of reactive nitrogen species (RNS) sources. (**A**) Events leading to oxidized and nitrated LDH. RNS (represented as NO_2_^−^/H_2_O_2_/ONOO^−^) can nitrate LDH, resulting in decreased V_max_. (**B**) Cardiac tissue was obtained from endothelial nitric oxide synthase (eNOS) and myeloperoxidase (MPO) knock-out (KO), and the respective wild-type (WT), mice. Basal nitration of LDH (p38^nt^) was analyzed by immunoblot, and immunosignals quantified densitometrically. Data are expressed as % of control (i.e., of the respective WT animals) and represent means ± SEM of n=3-5 animals. NO_2_-LDH depended on both eNOS and MPO

### Further MALDI-TOF mass spectrometry analysis of the nitration site identifies the nitrated residue as tryptophan and not tyrosine

A similarly frequent assumption about NO_2_Tyr immunosignals is specificity with respect to the nitrated amino acid. Other nitration target amino acids, however, have been reported, most prominently tryptophan (*28*–*30*). To identify the physiologically nitrated LDH residue(s), heart tissue was homogenized and subjected to 2D-PAGE in-gel digestion prior to MALDI-TOF MS analysis. Surprisingly, signals consistent with a nitrated tryptophan at position 324 of peptide 319-328 (SADTLWGIQK) of LDH (NO_2_Trp-LDH) were detected (Fig. 5A). We cannot exclude nitration of other peptides of LDH that were below our level of detection. Nevertheless, our results strongly point to NO_2_Trp being detected by the supposedly anti-NO_2_Tyr pAb1.

**Fig. 5.**
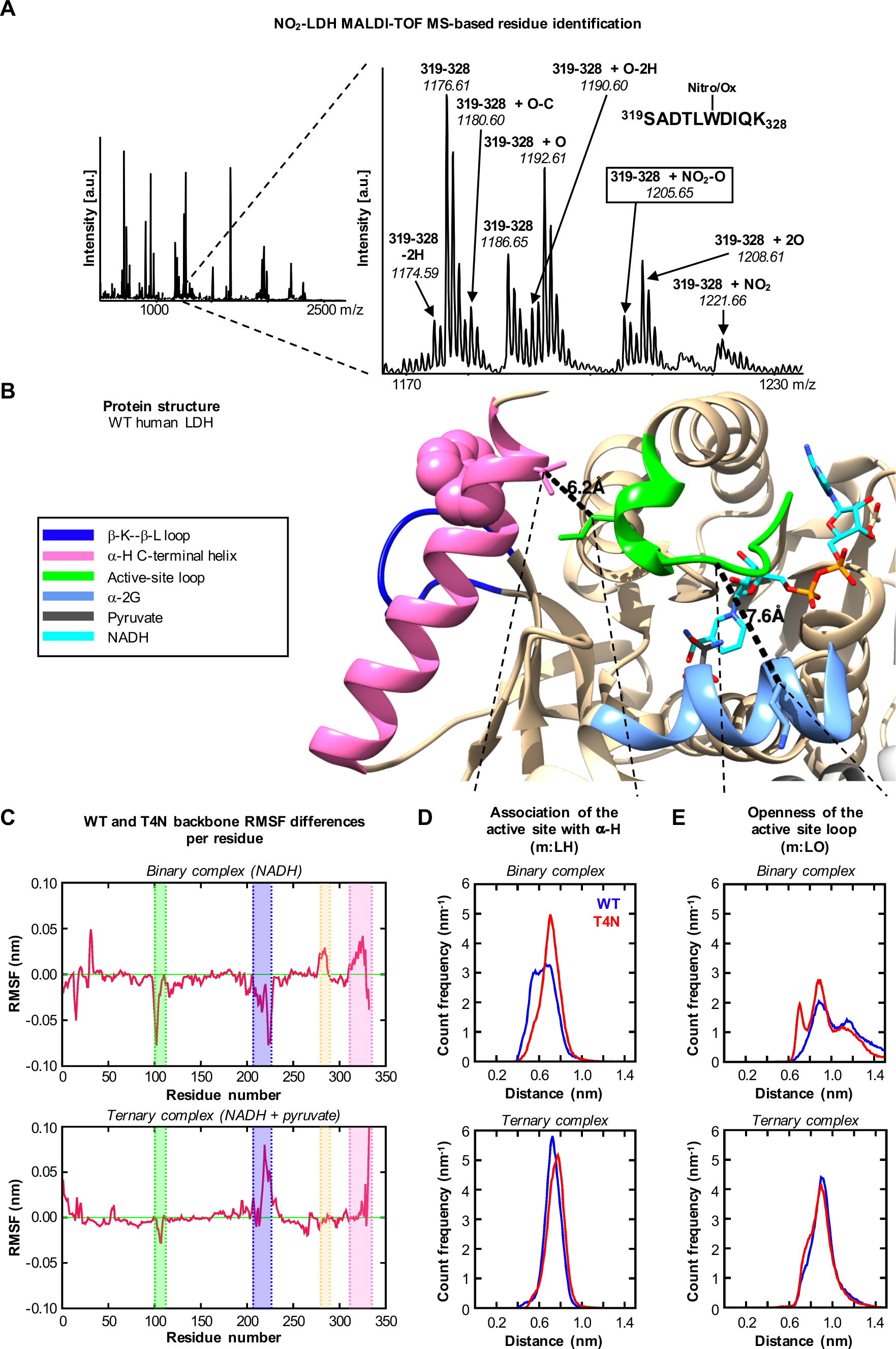
Identification of LDH Trp-324 as physiologically nitrated and oxidized. The interaction of the active site loop with the C-terminal α-helix is essential and restricted in NO_2_Trp-LDH. (**A**) Rat heart homogenate was subjected to 2D-PAGE and Coomasie staining; spots were subjected to in-gel digestion and MALDI-TOF MS analysis. Leftft panel: Mass spectrum derived from a spot corresponding to rat LDH over the range m/z 600 to 2700. Right panel: An expansion of the same spectrum over the range m/z 1170 to 1230, showing peptide ions assigned primarily to the peptide 319-328 (319SADTLWDIQK328), and its oxidized and nitrated derivatives. Major peptide ions are labeled with their observed m/z values and corresponding LDH mature protein amino acid intervals, as assigned by Mascot™, and additionally by manual comparison to predicted values calculated from the LDH sequence and oxidative and nitrative modifications. The ions correspond to the peptide 319-328 Trp (W)-nitrated species (319-328 + NO2) and its typical high energy decomposition product (319-328 + NO_2_-O), as well as ions corresponding to the characteristic family of Trp oxidation forms, hydroxytryptophan (319-328 + Ox), didehydro-hydroxytryptophan (319-328 + Ox-2H), dihydroxytryptophan (319-328 + 2Ox), and kynurenin (319-328 + Ox-C). Other peptide ions display single (+ Ox) and double (+ 2Ox) oxidation, or didehydrogenation (− 2H). (**B**) Distance metrics mapped onto the crystal structure of WT (1I0Z). Color scheme: β-K--β-K loop, dark blue; α-helix, pink; active-site loop, green; α-2G, light blue; pyruvate, grey; NADH, cyan. Residue Trp-324 is shown in space filling. (**C**) Plot of the differences between T4N and WT backbone RMSF per residue (T4N – WT) in the binary complex. Structures are highlighted using the same color scheme as previously described plus the juxtaposed loop to the C-terminal α-helix which is highlighted in light orange. (**D**, **E**) Histograms of the distribution of distances. Blue WT; Red T4N. Metrics m:LH (107:CB-326:CB) showing the juxtaposition of the active-site loop and α-C-terminal helix and m:LO (103:CA-243:CB) showing the open/closed state of the active-site loop.

### NO_2_Trp-LDH molecular dynamics simulations suggested higher restriction of the activity-essential interaction of the active site loop with the C-terminal α-helix

Since we did observe a V_max_ effect on NO_2_-LDH, we wanted to investigate whether there is a likely structural correlate for this via Trp-324. We therefore modeled the nitration into the structure of tetrameric and binary human LDH, and then performed molecular dynamics simulations. For distance metrics of human LDH and key structural components refer to Fig. 5B. The root-mean squared fluctuations (RMSF) of all atoms in each residue were averaged across the four subunits in each combined 3.5 μs trajectory (Fig. S3). Subtracting the wildtype tetramer (WT) RMSF from the T4N (tetramer with NO_2_Trp-324 residue on each subunit) highlighted differences in mobility between the dynamics of the WT compared with NO_2_Trp-LDH (Fig. 5C). Positive values in the resulting graphs indicated greater mobility of residues in T4N compared with WT and *vice versa*. Moreover, distance histograms (Fig. 5, D and E) show an increased mobility in the binary complexes compared with the ternary, expressed as a broadening of the histograms.

We found that NO_2_Trp-LDH weakened the interaction between α-H and the active site loop. This decoupling led to a greater population of closed and “over-closed” states of the T4N active-site loop. By corollary, in WT LDH, the open active-site loop states are partly stabilized by interaction with the α-H helix. Therefore, nitration of LDH Trp-324 is expected to decrease LDH activity, consistent with our enzyme-kinetic findings (Fig. 2, D and E).

## Discussion

We here close three major knowledge gaps in our understanding of protein nitration: specificity, origin, and disease relevance. With respect to the specificity of protein nitration, a vast heterogeneity of proteins are detected by supposedly uniform anti-NO_2_Tyr antibodies. Moreover, tryptophan nitration (NO_2_Trp) was a cofounding factor of our apparent NO_2_Tyr-immunosignals. Protein NO_2_Trp formation has been described in isolated proteins treated with H_2_O_2_/NO_2_^−^ *in vitro* (*31*–*34*), peroxynitrite-exposed cells (*35*), and as physiological modification during aging in rat heart mitochondria (*29*). Trp is an alternative target of nitration that deserves more attention (*36*, *37*) and immunosignals by anti-NO_2_Tyr antibodies should rather be interpreted as NO_2_Tyr/ NO_2_Trp signals unless purified and chemically validated. Rebrin et al. (*29*) reported Trp nitration of succinyl-CoA:3-ketoacid CoA transferase (SCOT), which could not be reproduced by Wang et al. (*38*). Whether Trp nitration has a functional consequence on SCOT remained unclear because both activation and inactivation were observed (*29*). In hindsight, SCOT nitration is also the first documented example where Trp nitration has been mistaken as Tyr nitration based on cross-reactivity of a supposedly NO_2_Tyr specific antibody (*39*), and many more of these examples may exist in the literature.

With respect to the origin of the nitrating nitrogen in NO_2_Tyr/ NO_2_Trp positive proteins, we show that both eNOS/NO and MPO/nitrite pathways, and possibly others, need to be equally considered and that the *a priori* assumption that NO_2_Tyr immunosignals indicate NO scavenging by superoxide is not valid. Nevertheless, the effects of KO of eNOS and MPO are significant but not great. Previous data shows that NO modifications remains present when NO synthesis and dietary nitrite/nitrate intake (e.g., in eNOS^−/−^ animals on a low nitrite/nitrate diet and under the acute action of L-NIO, a NOS-inhibitor) is suppressed (*40*). Moreover, NO levels remain essentially unchanged upon deletion of each NOS isoform, suggesting a compensatory mechanism maintains basal levels of NO metabolites in tissues.

With respect to disease relevance, probably our most relevant finding is that protein tryptophan nitration is a physiological mechanism modulating an enzyme, cardiac LDH, activity. This represents a similar dogma shift as for ROS formation, which also should not be *a priori* considered as oxidative stress and a disease marker. Thus, both ROS and reactive nitrogen and protein nitration represent physiological events, even though they may be upregulated in disease as we show for diabetes and cardiac stress (Fig. 3).

Not many specific functional consequences have been described for protein nitration. Assuming Trp nitration and activity modulation of both LDH (decreasing activity) and SCOT (increasing activity) would occur, this may be a protective mechanism, allowing the heart to better utilize ketone metabolism for energy production at a time when other metabolic processes may be diminished (*28*, *29*). Molecular dynamics simulations were performed on NO_2_-LDH, aiming to understand the impairment of the enzyme activity (Fig. 5). While differences in activity between WT and T4N of a factor of two is significant in physiological terms, it corresponds to minor changes in the free energy states in the catalytic process. The catalytic cycle of LDH is a well understood ordered bi-bi reaction mechanism where substrate and product enter and leave the active-site, with concomitant closing and opening of the active-site loop, after cofactor binding. Detailed kinetic characterization of LDH from *Bacillus stearothermophilus* (*41*) and *Plasmodium falciparum* (*42*) has revealed this loop movement to be a rate limiting step in the reaction mechanism. Any perturbation of this delicately balanced conformational change is expected to compromise the catalytic activity of the enzyme. We suggest that conformational movement of the active-site loop in cardiac LDH is also a rate limiting step and the weakening of the interaction between this loop and the α-H helix in T4N (as described in our results) is responsible for the impaired catalytic activity. Whether tryptophan nitration effects the conversion of either pyruvate or other 2-oxo/-2-hydroxy acids warrants further investigation.

Like many enzymes, LDH post-transcriptional activity is regulated by phosphorylation and acetylation of amino acid residues. The tyrosine kinase FGFR1 has been shown to directly phosphorylate LDH at Y10 and Y83 (*43*). Y10 phosphorylation of LDH promotes active, tetrameric LDHA formation, whereas phosphorylation of Y83 promotes NADH substrate binding. Our finding obviously promotes the opposite, suggesting an antagonistic effects of tyrosine phosphorylation and tryptophan nitration. Inhibition of LDH reduces aerobic glycolysis, gene transcription, channel complex regulation, molecular chaperones and cell cycle regulation. Cells are then forced to use oxidative phosphorylation and pyruvate enters the mitochondria. This leads to reactive oxygen species (ROS) generation (*44*). Since the nitration is already a ROS-induced event, this may be a mechanism of ROS-induced ROS formation (*45*), which in heart is protective triggering for example capillary density and angiogenesis (*46*, *47*).

In conclusion, cardiac LDH appears to be the first protein basally regulated by nitration and not only a loss-of-function biomarker of metabolic stress. NO_2_-LDH has a lower V_max_ and the nitration is further increased under cardiac metabolic stress. Furthermore, *in vivo* NO_2_Tyr immunoreactivity not necessarily reflects Tyr nitration or solely NO scavenging. Instead, it may indicate nitrite/MPO-dependent Trp nitration and physiological signaling. This has wide implications on re-interpreting previous data using NO_2_Tyr immunoreactivity as a biomarker and on our understanding of nitration as a physiological protein modification and signaling mechanism.

## Materials and Methods

### Chemicals

Immunoblotting reagents and column materials were obtained from Amersham Pharmacia Biotech (Freiburg, Germany); membranes, from Bio-Rad Laboratories (Hercules, CA, USA). Primary antibodies against nitrotyrosine were generous gifts from L.O. Uttenthal and J.S. Beckman or purchased from HyCult Biotechnology (Uden, Netherlands), Calbiochem-Novabiochem GmbH/Merck Millipore (Bad Soden/Darmstadt, Germany), Alexis/Axxora GmbH (Grünberg/Lörrach, Germany) and Upstate Inc./Millipore (Lake Placid, NY, USA/Darmstadt, Germany). As secondary antibodies, we used horseradish peroxidase (HRP)-conjugated goat anti-rabbit immunoglobulins for polyclonal antibodies or anti-mouse immunoglobulins for monoclonal primary antibodies from DAKO Diagnostika GmbH/Agilent Technologies (Hamburg, Germany). For gel loading we used Rotiload buffer from Carl Roth GmbH + Co. KG (Karlsruhe, Germany). The Oxyblot kit was supplied by Intergen Company/Millipore (Purchase, NY, USA/Darmstadt, Germany). Nonfat dry milk was obtained from Nestlé Inc. Carnation (Solon, Ohio, USA); Cell culture medium M199, from PAA Laboratories GmbH (Martinsried/Cölbe, Germany); Nitrating reagent, H_2_O_2_/NO_2_^−^, from Alexis/Axxora GmbH; Complete Protease Inhibitor Cocktail, from Roche Diagnostics GmbH (Mannheim, Germany). Porcine heart LDH was purchased from MyBiosource. All other chemicals were of the highest purity grade available and purchased from either Sigma-Aldrich Chemie GmbH (Taufkirchen, Germany) or Merck KG (Darmstadt, Germany).

### Animals

Adult male Wistar Unilever rats (270-320 g) were purchased from Harlan-Winkelmann (Borchen, Germany). eNOS and MPO KO mice were generated and bred as described previously (*20*). Rats and mice were housed under standard or SPF conditions, respectively, at a temperature of 18-20°C and a day-night rhythm of 12 h, with access to water and food ad libitum. Porcine hearts were obtained from a local abattoir.

### Nitration of proteins by peroxynitrite

If not stated otherwise, nitration was performed by continuous mixing protein and nitrating reagent (H_2_O_2_/NO_2_^−^) in a 1:5 molar ratio in phosphate buffer pH 7.3 (15 mM KH_2_PO_4_, 80 mM Na_2_HPO_4_) for 5 min at room temperature. Due to the strong alkaline storage conditions of H_2_O_2_/NO_2_^−^, the pH in the samples was immediately readjusted to neutral pH with HCl. No further purification or dialysis steps were conducted. For longer storage, samples were aliquoted and frozen at −20°C and used within 1 month or mixed with gel loading buffer and boiled for 10 min at 95°C for daily use. In this case, no change in signal characteristics or intensities was observed over time.

### SDS-PAGE, western and dot-blot analysis

Protein samples were separated by sodium dodecylsulfate-polyacrylamide gel electrophoresis (SDS-PAGE) under reducing conditions using 10% polyacrylamide gels. Alternatively, pre-cast gels (NuPAGE Novex 10% bis-Tris Midi Gel 1,0 mm × 26 well, Invitrogen) were used according to the manufacturer’s protocol with MES running buffer XCell SureLock Midi-Cell running tanks. For mass spectrometry proteins were stained with colloidal Coomassie blue. Proteins were immobilized on nitrocellulose membranes using standard dot-blot or semi-dry blotting techniques (*21*). Efficiency of protein immobilization was evaluated by staining the membrane in 0.5 % (w/v) Ponceau S in 3 % (w/v) trichloroacetic acid (TCA). Membranes were treated with 5 % (w/v) nonfat dry milk in Tris-buffered saline (TBS, 20 mM Tris/HCl pH 7.5, 150 mM NaCl) containing 0.1 % (v/v) Tween 20 (TBS-T) for 1 hour to block nonspecific protein binding. Then the membrane was incubated overnight at 4°C with the first antibody in 5% (w/v) nonfat dry milk in TBS-T and afterwards washed 3 × 10 min with TBS-T. To test the specificity for NO_2_Tyr, antibodies were pre-incubated with 3 mM NO_2_Tyr ^(footnote1)^. Alternatively, in order to reduce NO_2_ groups of nitrated proteins to NH_2_ groups, blots were first incubated with the reducing agent dithionite (1 mM) in 100 mM sodium borate pH 10 for 5 min at room temperature, washed 3 × 10 min with TBS-T and then incubated with anti-NO_2_Tyr antibody (*48*). As second antibody, we used horseradish peroxidase (HRP)-coupled antibodies against rabbit immunoglobulins (Ig) for polyclonal first antibodies or against mouse Ig for monoclonal first antibodies. HRP-labeled conjugates against the first antibodies were dissolved in 5% (w/v) nonfat dry milk in TBS-T. After washing 3 × 10 min with TBS-T the immunosignals were developed using the ECL detection reagent by Amersham Biosciences and blots were analyzed using a Kodak chemiluminescence Imager (Scientific Imaging System, Eastman Kodak Company, New Haven, USA). The resolution of the imager was 752 × 582 pixels and the linear area of signal quantification was verified by repeated measurements using increasing protein amounts. Light signals were directly quantified employing the Kodak 1D Image analysis software (version 3.5.2).

### Screening of tissues and determination of nitrated proteins

Several tissues from rat and pig were homogenized in 10 volumes (w/v) of lysis buffer, containing 25 mM triethanolamine HCl (TEA) pH 7, 1 mM EDTA, 5 mM dithiothreitol, 50 mM NaCl, 10 % (v/v) glycerol and Complete™ Protease Inhibitor Cocktail (Roche, Mannheim, Germany). For long-term storage, the homogenates were shock frozen in liquid nitrogen and stored at −20 °C. For daily use, gel loading buffer was added to the samples and heated to 95°C for 10 min. Any remaining particles were pelleted by centrifugation in a desktop centrifuge (10000 × *g*, 10 min, 4°C). The samples were first screened by use of all antibodies listed in Tab. 1 by dot blot analysis. Tissues that yielded more than 4 hits by all antibodies were subjected to further analysis by SDS-PAGE and western blot as described above.

### Characterization and isolation of a 38 kDa protein in pig heart tissue

A column packed with 2 ml of Blue Sepharose resin (Amersham Bioscience) was washed and pre-equilibrated with 10 mL lysis buffer. Pig heart supernatant (1 mL) was applied to the column. The flow-through was subsequently applied to 1 mL Q-Sepharose column that was also washed and equilibrated with lysis buffer. This column was then washed again with lysis buffer (5 mL) and the bound proteins were eluted with 2 mL of lysis buffer containing 1 M NaCl. The partially purified 38-kDa protein was mixed with gel loading buffer, boiled at 95°C for 10 min and subjected to SDS-PAGE. It was stained with colloidal Coomassie blue (*23*). The stained protein band at 38 kDa was cut out, destained and enzymatically digested. Tryptic fragments were separated by high pressure liquid chromatography (HPLC). Fractions containing tryptic peptides were subjected to matrix-assisted laser-desorption/ionization mass spectrometry (MALDI-MS, Bruker Reflex III™, Bruker Daltonics, Bremen, Germany). Peptide sequences were compared with proteins in the NCBIprot database.

### 2D-PAGE analysis of rat heart tissue

2D-PAGE analysis was conducted using standard conditions (*49*). Briefly, 7 cm ReadyStrips™ IPG strips (BioRad) of pI 3-10 were subjected to active rehydration overnight with heart homogenate in Destreak™ IEF buffer, supplemented with 0.02% pI 3-10 ampholites (BioRad). Isoelectric focusing was performed in a Protean™ IEF cell (Bio-Rad) using a ramped voltage up to 4000 V, until a total of approximately 10,000 Vh had elapsed. Strips were subjected to reduction, alkylation, and equilibration for the second dimension, using Equilibration Buffers I & II (BioRad) for 30 min each, then SDS-PAGE, using 10% polyacrylamide NuPage™ Novex Bis-Tris gels (Invitrogen). Gels were subsequently Coomassie stained (Gelcode™ Blue, Pierce).

### In-gel digestion of rat heart samples

In-gel digestion was conducted as previously described (*50*). Briefly, the well-resolved protein spot of interest was punch-excised using wide-orifice pipette tips. Gel pieces were destained with 100 mM ammonium bicarbonate, pH 9/50% acetonitrile, and were washed three times with wash solution 1 (100 mM ammonium bicarbonate, pH 9), wash solution 2 (100 mM ammonium bicarbonate, pH 9/50% acetonitrile), followed by wash solution 3 (100% acetonitrile). Gel pieces were swelled in 50 mM ammonium bicarbonate, pH 9/10% acetonitrile, containing approximately 10 ng Trypsin Gold (Promega) and incubated overnight at 37°C. Peptides were extracted (each time for 20 min) once with 20 mM ammonium bicarbonate, pH 9, twice with 1% trifluoroacetic acid/50% acetonitrile, followed by once with 100% acetonitrile. The entire extraction process was performed in duplicate, and all extraction supernatants were pooled and spun to dryness in a SpeedVac™ (Thermo-Savant, Waltham, MA).

### MALDI-TOF MS of rat heart samples

Peptides were desalted and prepared for MS using ZipTips™ (Millipore). Mass spectra were obtained after purified peptides were co-crystallized with the matrix 2,5-dihydroxybenzoic acid onto AnchorChip™ targets (Bruker Daltonics, Billerica, MA) using the dried-droplet technique (*29*, *51*) and a Reflex IV™ MALDI-TOF MS instrument (Bruker-Daltonics) in positive ion, reflectron mode, over the range 400-8000. The laser intensity was adjusted to be ~3% above the threshold value for peptide ion signals. Signals from 100 to 200 laser shots were summed for each mass spectrum. External and internal calibration was achieved to within 50 ppm using the masses of known purified peptide standards and trypsin autolysis peptides.

### MS data analysis and peptide mass fingerprinting

Peak lists from MALDI-TOF mass spectra (derived using MoverZ™ software, Genomic Solutions, Ann Arbor, MI) were submitted on-line to the Mascot™ search engine (Matrix Science, London, UK) for peptide mass fingerprinting (PMF) analysis against the SwissProt or NCBI non-redundant protein databases, using the following restrictions: a) Rattus sp., b) trypsin digestion with up to 3 missed cleavages, c) +/− 50 ppm error, and d) cysteine carbamidomethylation and methionine oxidation as fixed and variable modifications, respectively. LDH was unambiguously identified by PMF, with a Mascot™ Score of 150 (corresponding to an E value of 6 × 10^−12^, the likelihood of a false assignment) and a sequence coverage of 65% from 35 matched peptide ions, given search parameters. Additional peptide ion assignments (as well as validation of Mascot™ assignments) were made by manual inspection of spectra using theoretical digest and post-translational modification mass values for LDH.

### LDH activity, oxidation and nitration by H_2_O_2_/NO_2_^−^

Commercial LDH was treated with different amounts of peroxynitrite as described above. Aliquots of H_2_O_2_/NO_2_^−^ were diluted with 2 M NaOH to reach the desired concentrations indicated in **Fig. 2**. LDH activity and kinetic parameters of exogenously nitrated LDH were measured using a fluorescence-based assay, using excitation/emission setting of 340/470 nm, respectively. The assay used uses the fluorescent properties of NADH which can be excited at 340 nm and emission is measured at 440 nm. The oxidized form of NADH, NAD^+^, does not have these properties. Measurements were collected every 20 seconds during 5 minutes. The final concentrations in the reaction were LDH, 2 mU/mL, NADH 0.12 mM and Sodium pyruvate 1.13 mM. Nitration of LDH was measured as the intensity of the NO_2_Tyr immunosignal in arbitrary units.

### Molecular Dynamics Methods

#### System setup

Two simulation systems were setup based on the crystal structure of the ternary complex of human-heart LDH (pdb 1I0Z). One being the wild type tetramer (WT) and the other being the tetramer with a nitro group substituted at each Trp-324 residue (T4N). The oxamate molecules were converted to pyruvate and acpype (*52*) used to generate Amber GAFF parameters for the substrate and cofactor molecules. A combination of GAFF parameters for nitrotryptophan and analogy was used to generate a residue library entry for a nitrotryptophan in the Gromacs AmberSB99-ildn forcefield (*53*). Gromacs tools were used to add hydrogen atoms to all residues consistent with pH 7 and to protonate the catalytic histidine (His-193). The complexes were solvated in boxes of TIP3P water containing 0.15 M sodium chloride ions. Calculations were performed at NTP under period boundary conditions and the PME method used for long range electrostatics. The pressure was 1 Barr and the temperature 300 K for all molecular dynamics (MD) simulations. Both systems were relaxed by 2000 steps of steepest descent energy minimisation followed by 0.2 ns MD while restraining the protein atoms to their initial positions.

Another pair of simulation systems were prepared in the same fashion, with the pyruvate removed. This gave a total of four simulation systems, referred to as WT-ternary, WT-binary, T4N-ternary and T4N-binary.

#### Trajectory development and analysis

Three 200 ns simulations (initialized with different velocities) were performed for each of the four simulation systems. These simulations were performed on the University of Bristol HPC machines BlueCrystal and BlueGem. Structures were extracted every 5 ns from each trajectory and energy minimized. This gave 120 starting structures for each of the four systems for further parallel trajectory development. Each new trajectory was initialized with a fresh set of random velocities and run for 28-31 ns on the UK National HPC machine Archer, giving around 3.5 μs combined simulation data for each system. All simulations were run and analyzed using the Gromacs 5.1.2 suite of software (*54*) and visualizations performed with VMD 1.9.2 (*55*) and Chimera 1.11 (*56*).

### Molecular Dynamics Analysis

The active site loop (residues 98-110) was found to be less mobile in T4N than WT in the binary complex (LDH+NADH), while the C-terminal helix (α-H) and the juxtaposed loop (residues 277-287) were more mobile in T4N. The lower mobility of the active site loop is also detectable in the ternary complex (LDH+NADH+pyruvate) albeit greatly attenuated, while the C-terminal helix mobility is similar to the WT. The other main features in these plots are the differences in mobility of the 205-225 loop. However, as this is one of the most flexible parts of the protein and is not located near the active site we do not consider these differences to impact the activity of the two proteins. By contrast, the C-terminal side of the active site loop juxtaposes the C-terminal end of the final helix in the protein, α-H (residues 309-332). A crucial contact between these two structural elements is made by Leu-107 and Ile-326. Hence, we define one metric reporting the association of the active site loop with α-H via the distance between CB atoms of these residues, as m:LH. Likewise, two residues spanning the mouth of the active site Gly-103 and Lys-243 were chosen to monitor openness of the active site loop, or m:LO. Distance histograms show the distribution of these distances over each of the four 3.5 μs combined trajectories. They indicate an increased mobility in the binary complexes compared with the ternary, expressed as a broadening of the histograms.

The distance between the active site loop and α-H (m:LH) is slightly shorter on average for WT than T4N in the ternary complex and this difference is amplified in the binary complex where about half of the WT loops interact more tightly with α-H than T4N. Comparison with the second metric monitoring active site loop closure (m:LO) in the binary complex shows that the T4N loop spends more time at a distance comparable with the ternary complex than the WT and also is an “over-closed” state. The results of the ternary simulations are again more similar, but a tendency towards and “over-closed” state persists.

### Synthesis and Characterization of Nitrotryptophan

Nitrotryptophan (NO_2_Trp) was synthesized according to Moriya, et al. (*15*). 20 g L-Trp and 0.25 g urea were suspended in 350 mL glacial acetic acid. Under vigorous stirring at 4°C, 10 mL 24 M nitric acid in 30 mL glacial acetic acid were added. After the suspension changed to a yellow clean solution, it was stirred at 4°C until it changed again to a suspension. Then an additional 10 mL fuming nitric acid in 30 mL glacial acetic acid were added drop by drop at 15°C. After stirring overnight at room temperature, the resulting yellow precipitate was collected by filtration and washed with glacial acetic acid. The crude product was recrystallized in 100 mL 2 M nitric acid at temperatures below 50°C and converted to the free amino acid in 100 mL hot water by the addition of sodium carbonate until the pH was neutral. As analyzed by its infrared (IR) and nuclear magnetic resonance (NMR) spectra, no 4-NO_2_Trp was detectable. Because 7-NO_2_Trp cannot be synthesized by the action of 24 M nitric acid on the tryptophan ring (*16*), the products of the synthesis with respect to UV spectrum are a mixture of 5/6-NO_2_Trp. NO_2_Trp crystallized as yellow needles with a melting point at 265°C (decomp.). The compound was characterized according to: UV (in 1 N NaOH) 330 nm, 380 nm; IR (KBr) (−NH_3_^+^) 3190 cm^−1^, *v*(C=C) and amide I 1600-1650 cm^−1^, amide II and δ(−NH_3_^+^) 1550-1600 cm^−1^, *v*_as_(−NO_2_) 1495 cm^−1^, *v*_s_(−NO_2_) 1330 cm^−1^, predominantly 5/6-NO_2_ substitution at 818 cm^−1^ instead of 4-NO_2_ substitution at 780 cm^−1^; NMR (D_2_O-NaOH, δ 4.7 HDO) δ 2.9−3.1 (2H, m), 3.4−3.6 (1H, m), 7.35 (1H,s), 7.5 (1H, d, J= 9 Hz), 7.7 (1H, dd, J= 2 and 9 Hz), 7.95 (1H, d, J= 2 Hz).

### Animal models of disease

To examine LDH nitration under pathophysiological conditions, two *in vivo* models involving nitrative stress were assessed, including a short- and longer-term streptozotocin diabetes model in the rat, and a murine model of doxorubicin-induced cardiotoxicity.

### Streptozotocin model of diabetes mellitus

At an age of 10 weeks, rats (n=12) received a single intraperitoneal injection of 60 mg/kg BW of streptozotocin (STZ) dissolved in citrate buffer 50mM pH 4,5. Age matched control rats (n=12) received vehicle injections with citrate buffer. Hyperglycemia was defined as a positive test of glucose in urine (Haemo-Glukotest, Boehringer Mannheim, Germany) and a non-fasting blood glucose level higher than 11 mmol/L (hexokinase method) one week after STZ application. Rats were sacrificed 3 (n=12) and 16 weeks (n=12) after diabetes induction by CO_2_-inhalation. After thoracotomy, hearts were excised and freed of connective tissue while immersed in modified Krebs-Hepes buffer pH 7.35 containing 99 mM NaCl; 4.7 mM KCl; 1.9 mM CaCl_2_; 1.2 mM MgSO_4_; 25 mM NaHCO_3_; 1 mM K_2_HPO_4_; 20 mM Na-Hepes; 11.1 mM D-glucose. Hearts were shock frozen in liquid nitrogen, pulverized, and homogenized in lysis buffer. For SDS-PAGE and western blot, gel loading buffer was added and protein concentrations were determined after TCA precipitation by the method of Lowry.

### Doxorubicin cardiotoxicity model

Doxorubicin (DOX, Adriamycin) is a broad-spectrum anthracycline antibiotic that is used to treat a variety of cancers. However, the clinical use of DOX is limited because of its serious cardiotoxicity, possibly involving increased oxidative stress, alteration of cardiac energetic status and direct effects on DNA. Hearts from DOX-treated mice were obtained as described elsewhere (*57*), and homogenized and shock-frozen as described above.

### Endogenous LDH nitration in eNOS-KO and MPO-KO mice

Hearts from eNOS-KO and MPO-KO mice were excised, washed in perfusion buffer, blotted dry on filter paper, weighed, and homogenized immediately in ice-cold 50 mM PBS, pH= 7.4, containing 2.5 mM EDTA, 1 mM DTT, 1% NP-40, phosphatase inhibitors (10 mM sodium fluoride, 10 mM sodium pyrophosphate, 2 mM sodium orthovanadate, and 50 mM β-glycerophosphate), and Complete™ protease inhibitor cocktail (Roche Diagnostics, Indianapolis, IN). Homogenates were incubated on ice for 30 min to facilitate solubilization, after which they were clarified by centrifugation at 12000 × g for 15 min at 4°C, aliquoted and stored at −80°C. Prior to use, samples were thawed on ice and assayed for protein concentration using the BCA method.

## Supplementary Materials

**Table S1.** Commercial and non-commercial antibodies detecting reactive nitrogen species-induced protein modifications.

**Fig. S1.** Anti-NO_2_-Tyr antibodies greatly vary in sensitivity and specificity in native tissues.

**Fig. S2.** Nitration and oxidation of LDH *in vitro* inversely correlates with LDH activity.

**Fig. S3.** Plots of the backbone RMSF per residue averaged over all subunits of all 120 simulations.

## Acknowledgments

The authors would like to thank G. Jansen for technical help, and Dr. P.M.H. Schiffers for carefully reading earlier stages of the manuscript. We also thank the Advanced Computing Research Centre at Bristol and BrisSynBio for access to the HPC machines BlueCrystal and BlueGem. We thank the HECBiosim Consortium for access to the UK National HPC facility Archer.

## Author contributions

H.H.H.W.S. designed research; J.F., D.T., T.N., C.I.A., A.S., P.B., C.S., V.S., P.R., A.J., A.G., D.H.P., R.B.S. and M.F. performed research; A.S., A.G., A.K., S.B., D.H.P., M.F. and H.H.H.W.S. contributed new reagents/analytic tools; J.F., D.T., D.H.P., and H.H.H.W.S. analyzed data; and C.N., J.F., C.S., D.H.P., M.F., A.I.C. and H.H.H.W.S. wrote the paper.

## Data and materials availability

Data and materials available upon reasonable request.

## Supplementary Materials

**Supplementary Table 1.** Commercial and non-commercial antibodies detecting reactive nitrogen species-induced protein modifications. Poly (pAb)- and monoclonal (mAb) antibodies from the indicated species were obtained from either commercial sources or kindly provided by the listed researchers. All antibodies, except pAb3 and mAb6, were raised against keyhole limpet hemocyanin (KLH) treated with ‘nitration reagent’, i.e. H_2_O_2_/NO_2_^−^. For pAb3, the hapten was nitrosocysteine (NOCys); for mAb6, nitrosotryptophan (NOTrp), and in both cases glutaraldehyde-coupled to KLH. Antibodies mAb1, mAb2 and mAb3 behaved identically and referred to the same reference (34), and were thus considered identical. Antibodies mAb4, mAb5 and pAb3 were later omitted from the study because of low titer and/or low affinity, which allowed to detect only highly abundant antigens such as H_2_O_2_/NO_2_^−^-treated bovine serum albumin (BSA), but not lower abundance endogenously nitrated proteins in native tissues. Antibody mAb6 was also excluded from further use because it reacted similarly with untreated and H_2_O_2_/NO_2_^−^-treated BSA. Consequently, we pursued the further use of the first three antibodies, pAb1, pAb2 and mAb1, for the present study (see Fig. 2).

| Abbreviation | Specificity | Antigen | Species | Source | Reference |
| --- | --- | --- | --- | --- | --- |
| pAb1 | anti-NO<sub>2</sub>Tyr | ONOO-KLH | rabbit | L.O. Uttenthal | (33) |
| pAb2 |  |  |  | J.S. Beckman | (34) |
| mAb1 |  |  | mouse |  | (34) |
| Identical to mAb1: |  |  |  |  |  |
| mAb2 | anti-NO<sub>2</sub>Tyr | ONOO-KLH | mouse | Alexis | (34) |
| mAb3 |  |  |  | Upstate | (34) |
| Rejected: |  |  |  |  |  |
| mAb4 | anti-NO<sub>2</sub>Tyr | ONOO-KLH | mouse | Calbiochem | (51) |
| mAb5 |  |  |  | HyCult biotechnology | (37) |
| pAb3 | anti-NOTrp | NOTrp-KLH | rabbit | Calbiochem | None |
| mAb6 | anti-NOCys | NOCys | mouse | Biolabs | None |

**Supplementary Figure 1.**
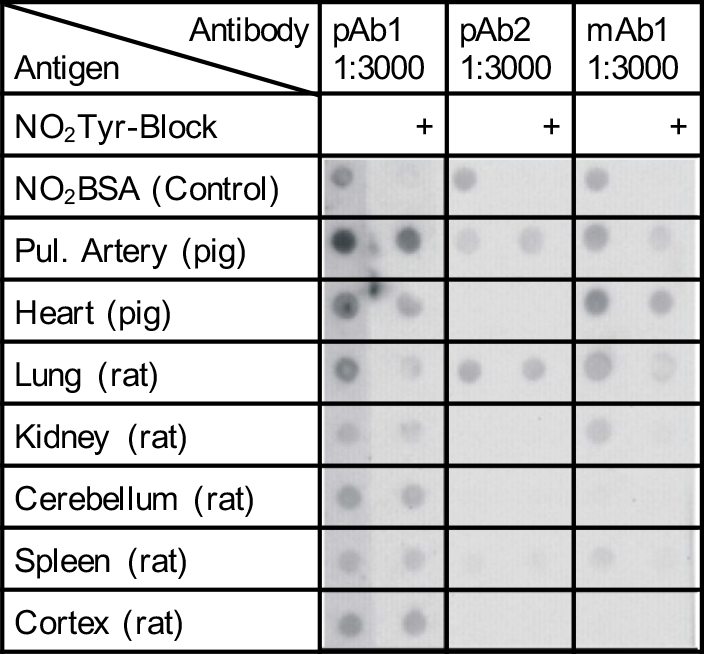
Anti-NO2-Tyr antibodies greatly vary in sensitivity and specificity in native tissues. Dot blot screen of tissue homogenates of porcine pulmonary artery and heart, and rat heart, lung, kidney, cerebellum, spleen and cortex. For each dot, 50 μg or, in the case of pulmonary artery, 100 μg of protein were blotted onto a nitrocellulose membrane. NO_2_Tyr-immuno-positive proteins were then detected using antibodies pAb1, pAb2 or mAb1. NO_2_BSA was used as a positive control; pre-incubation with 3 mM free NO_2_Tyr, as a negative control (+).

**Supplementary Figure 2.**
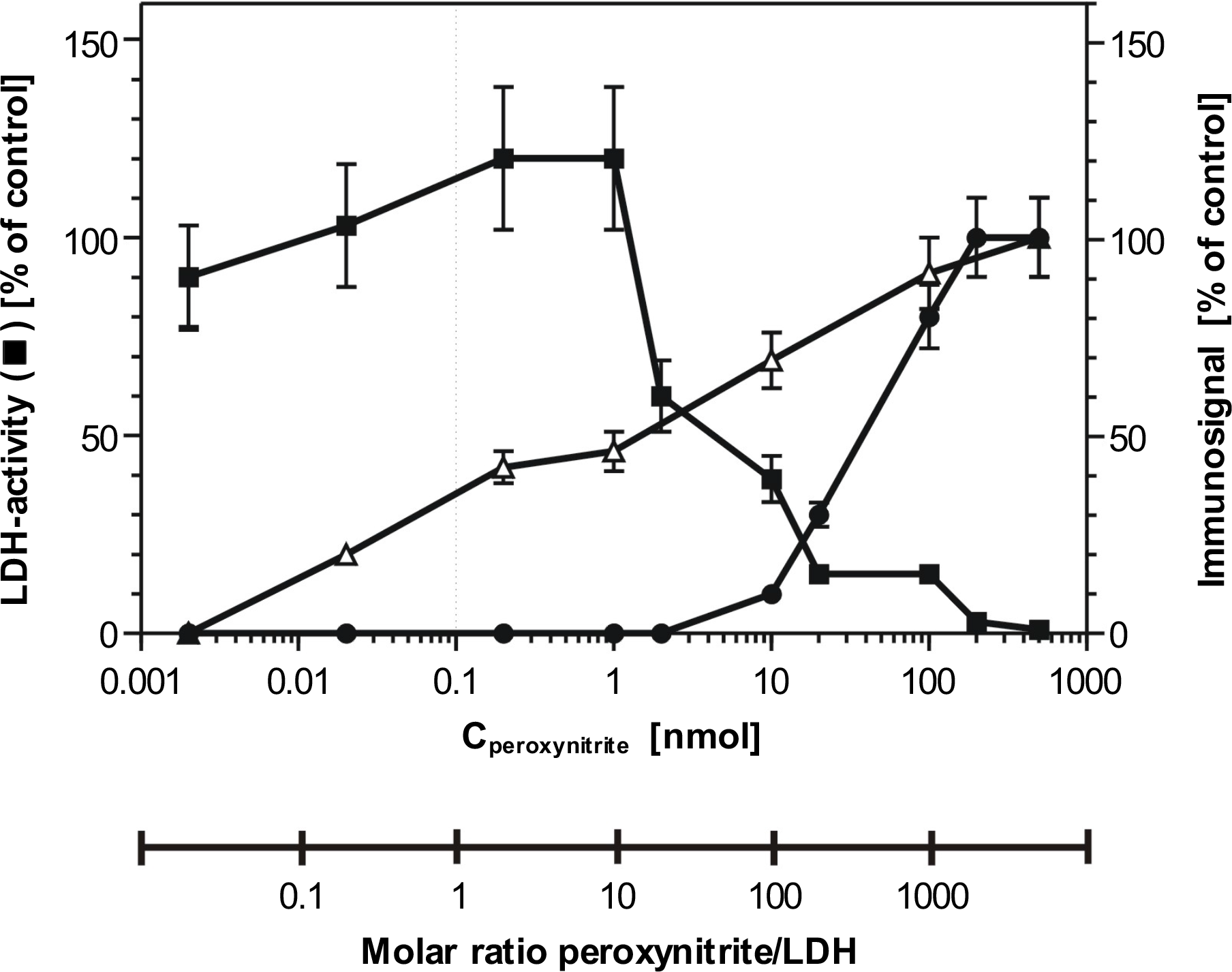
Nitration and oxidation of LDH *in vitro* inversely correlates with LDH activity. As LDH (0.1 nmol) was treated with increasing amounts of peroxynitrite, total LDH activity (∎, left axis) was determined by the NADH-coupled photometric assay of Warburg, as described, and all LDH samples were tested under the same conditions for the occurrence of nitration (⚫) using quantitative Western analysis with the NO_2_Tyr antibody pAb1 and oxidation (△) using the oxyblot detection kit (both, right axes).

**Supplementary Figure 3.**
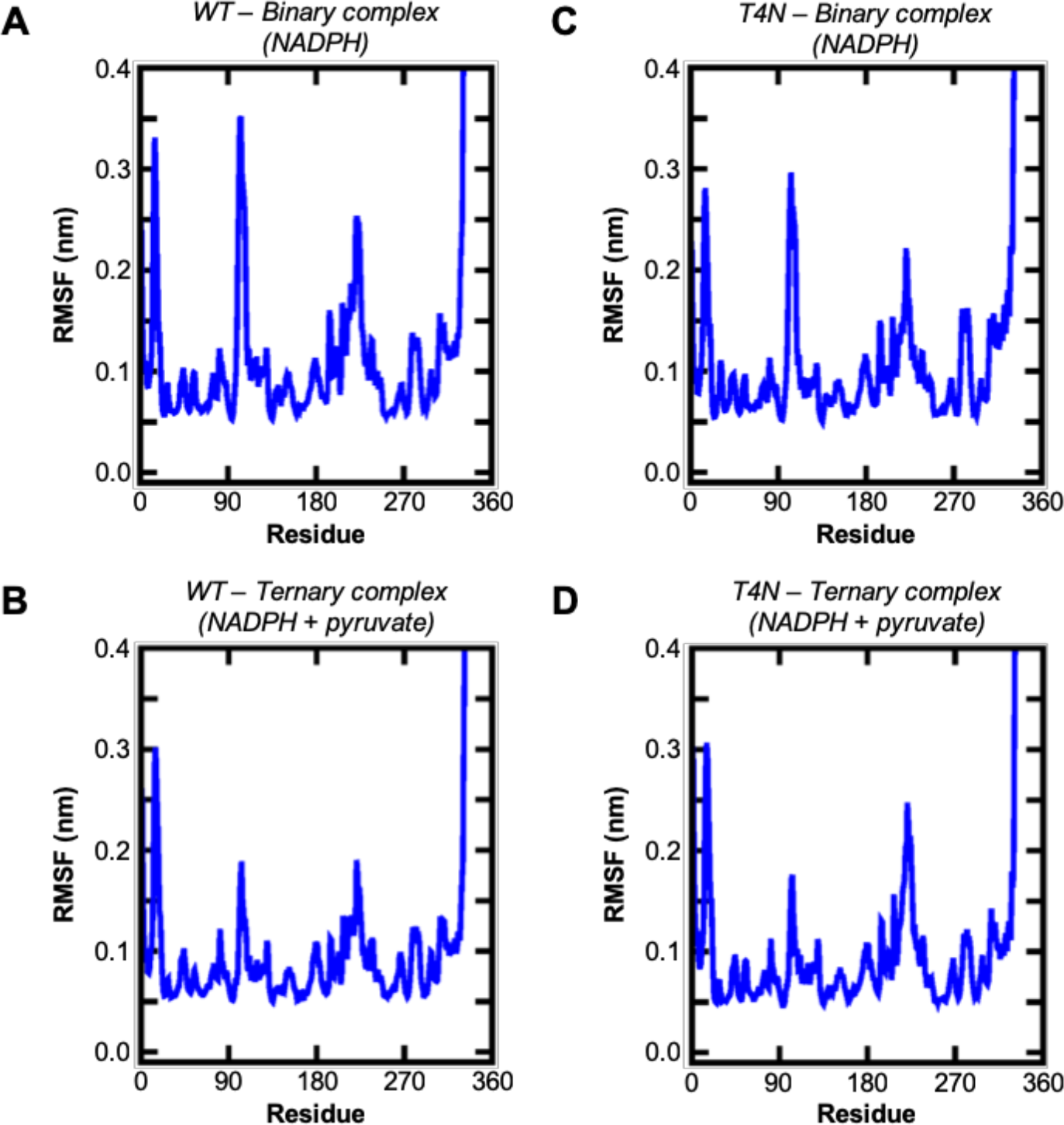
Plots of the backbone RMSF per residue averaged over all subunits of all 120 simulations. (**A**) WT-binary. (**B**) WT-ternary. (**C**) T4N-binary. (**D**) T4N-ternary.

## References and Notes

1. H. Sies, Oxidative stress: a concept in redox biology and medicine. Redox Biol. 4, 180–183 (2015).

2. J. S. Beckman, W. H. Koppenol, Nitric oxide, superoxide, and peroxynitrite: the good, the bad, and ugly. Am. J. Physiol. Physiol. 271, C1424–37 (1996).

3. R. Radi, Nitric oxide, oxidants, and protein tyrosine nitration. Proc. Natl. Acad. Sci. 101, 4003–4008 (2004).

4. M. E. Armitage, K. Wingler, H. H. H. W. Schmidt, M. La, Translating the oxidative stress hypothesis into the clinic: NOX versus NOS. J. Mol. Med. 87, 1071–1076 (2009).

5. S. Viappiani, R. Schulz, Detection of specific nitrotyrosine-modified proteins as a marker of oxidative stress in cardiovascular disease. AJP Hear. Circ. Physiol. 290, H2167–H2168 (2006).

6. M. H. Shishehbor, S. L. Hazen, Inflammatory and oxidative markers in atherosclerosis: Relationship to outcome. Curr. Atheroscler. Rep. 6, 243–50 (2004).

7. E. Ho, K. Karimi Galougahi, C.-C. Liu, R. Bhindi, G. A. Figtree, Biological markers of oxidative stress: Applications to cardiovascular research and practice. Redox Biol. 1, 483–491 (2013).

8. E. Fernández, J.-M. García-Moreno, A. Martín de Pablos, J. Chacón, May the Evaluation of Nitrosative Stress Through Selective Increase of 3-Nitrotyrosine Proteins Other Than Nitroalbumin and Dominant Tyrosine-125/136 Nitrosylation of Serum α-Synuclein Serve for Diagnosis of Sporadic Parkinson’s Disease? Antioxid. Redox Signal. 19, 912–918 (2013).

9. B. Halliwell, What nitrates tyrosine? Is nitrotyrosine specific as a biomarker of peroxynitrite formation in vivo? FEBS Lett. 411, 157–160 (1997).

10. Y. Z. Ye, M. Strong, Z.-Q. Q. Huang, J. S. Beckman, Antibodies that recognize nitrotyrosine. Methods Enzymol. 269, 201–209 (1996).

11. C. D. Reiter, R.-J. Teng, J. S. Beckman, Superoxide Reacts with Nitric Oxide to Nitrate Tyrosine at Physiological pH via Peroxynitrite. J. Biol. Chem. 275, 32460–32466 (2000).

12. N. Breusing, T. Grune, Biomarkers of protein oxidation from a chemical, biological and medical point of view. Exp. Gerontol. (2010), doi:10.1016/j.exger.2010.04.004.

13. J. B. Sampson, Y. Ye, H. Rosen, J. S. Beckman, Myeloperoxidase and Horseradish Peroxidase Catalyze Tyrosine Nitration in Proteins from Nitrite and Hydrogen Peroxide. Arch. Biochem. Biophys. 356, 207–213 (1998).

14. D. J. Bigelow, Nitrotyrosine-modified SERCA2: a cellular sensor of reactive nitrogen species. Pflügers Arch. - Eur. J. Physiol. 457, 701–710 (2009).

15. T. V. Knyushko, V. S. Sharov, T. D. Williams, C. Schöneich, D. J. Bigelow, 3-Nitrotyrosine Modification of SERCA2a in the Aging Heart: A Distinct Signature of the Cellular Redox Environment †. Biochemistry. 44, 13071–13081 (2005).

16. H. Li, D. D. Gutterman, N. J. Rusch, A. Bubolz, Y. Liu, Nitration and Functional Loss of Voltage-Gated K+ Channels in Rat Coronary Microvessels Exposed to High Glucose. Diabetes. 53, 2436–2442 (2004).

17. I. Y. Haddad, S. Zhu, H. Ischiropoulos, S. Matalon, Nitration of surfactant protein A results in decreased ability to aggregate lipids. Am. J. Physiol. Cell. Mol. Physiol. 270, L281–8 (1996).

18. G. Ferrer-Sueta et al., Biochemistry of Peroxynitrite and Protein Tyrosine Nitration. Chem. Rev. (2018), doi:10.1021/acs.chemrev.7b00568.

19. T. Akaike et al., Pathogenesis of influenza virus-induced pneumonia: involvement of both nitric oxide and oxygen radicals. Proc. Natl. Acad. Sci. 93, 2448–2453 (1996).

20. T. Akaike et al., 8-Nitroguanosine formation in viral pneumonia and its implication for pathogenesis. Proc. Natl. Acad. Sci. 100, 685–690 (2003).

21. T. J. Evans et al., Cytokine-treated human neutrophils contain inducible nitric oxide synthase that produces nitration of ingested bacteria. Proc. Natl. Acad. Sci. 93, 9553–9558 (1996).

22. T. Hayashi et al., Modulating role of estradiol on arginase II expression in hyperlipidemic rabbits as an atheroprotective mechanism. Proc. Natl. Acad. Sci. 103, 10485–10490 (2006).

23. C. Szabo et al., Protection against peroxynitrite-induced fibroblast injury and arthritis development by inhibition of poly(ADP-ribose) synthase. Proc. Natl. Acad. Sci. 95, 3867–3872 (1998).

24. M. L. Graham, J. L. Janecek, J. A. Kittredge, B. J. Hering, H. J. Schuurman, The streptozotocin-induced diabetic nude mouse model: Differences between animals from different sources. Comp. Med. (2011), doi:10.1152/ajprenal.00292.2015.

25. J. M. Berthiaume, K. B. Wallace, in Cell Biology and Toxicology (2007).

26. G. Peluffo, R. Radi, Biochemistry of protein tyrosine nitration in cardiovascular pathology. Cardiovasc. Res. 75, 291–302 (2007).

27. A. van der Vliet, J. P. Eiserich, B. Halliwell, C. E. Cross, Formation of Reactive Nitrogen Species during Peroxidase-catalyzed Oxidation of Nitrite. J. Biol. Chem. 272, 7617–7625 (1997).

28. T. Nuriel, A. Hansler, S. S. Gross, Protein nitrotryptophan: Formation, significance and identification. J. Proteomics. 74, 2300–2312 (2011).

29. I. Rebrin, C. Brégère, S. Kamzalov, T. K. Gallaher, R. S. Sohal, Nitration of Tryptophan 372 in Succinyl-CoA:3-Ketoacid CoA Transferase during Aging in Rat Heart Mitochondria †. Biochemistry. 46, 10130–10144 (2007).

30. J. M. Souza, G. Peluffo, R. Radi, Protein tyrosine nitration—Functional alteration or just a biomarker? Free Radic. Biol. Med. 45, 357–366 (2008).

31. T. Suzuki et al., Nitration and nitrosation of N-acetyl-l-tryptophan and tryptophan residues in proteins by various reactive nitrogen species. Free Radic. Biol. Med. 37, 671–681 (2004).

32. F. Yamakura, K. Ikeda, Modification of tryptophan and tryptophan residues in proteins by reactive nitrogen species. Nitric Oxide. 14, 152–161 (2006).

33. F. Yamakura et al., Nitrated and Oxidized Products of a Single Tryptophan Residue in Human Cu,Zn-Superoxide Dismutase Treated with Either Peroxynitrite-Carbon Dioxide or Myeloperoxidase-Hydrogen Peroxide-Nitrite. J. Biochem. 138, 57–69 (2005).

34. F. Yamakura, T. Matsumoto, H. Taka, T. Fujimura, K. Murayama, 6-Nitrotryptophan: a specific reaction product of tryptophan residue in human Cu, Zn-SOD treated with peroxynitrite. Adv. Exp. Med. Biol. 527, 745–9 (2003).

35. K. Ikeda et al., Detection of 6-nitrotryptophan in proteins by Western blot analysis and its application for peroxynitrite-treated PC12 cells. Nitric Oxide. 16, 18–28 (2007).

36. B. Alvarez et al., Peroxynitrite-Dependent Tryptophan Nitration. Chem. Res. Toxicol. 9, 390–396 (1996).

37. S. Herold, Nitrotyrosine, dityrosine, and nitrotryptophan formation from metmyoglobin, hydrogen peroxide, and nitrite. Free Radic. Biol. Med. 36, 565–579 (2004).

38. Y. Wang et al., The Nitrated Proteome in Heart Mitochondria of the db/db Mouse Model: Characterization of Nitrated Tyrosine Residues in SCOT. J. Proteome Res. 9, 4254–4263 (2010).

39. S. Marcondes, I. V Turko, F. Murad, Nitration of succinyl-CoA:3-oxoacid CoA-transferase in rats after endotoxin administration. Proc. Natl. Acad. Sci. 98, 7146–7151 (2001).

40. A. B. Milsom, B. O. Fernandez, M. F. Garcia-Saura, J. Rodriguez, M. Feelisch, Contributions of nitric oxide synthases, dietary nitrite/nitrate, and other sources to the formation of NO signaling products. Antioxid. Redox Signal. 17, 422–32 (2012).

41. H. M. Wilks et al., Designs for a broad substrate specificity keto acid dehydrogenase. Biochemistry. 29, 8587–8591 (1990).

42. D. K. Shoemark, M. J. Cliff, R. B. Sessions, A. R. Clarke, Enzymatic properties of the lactate dehydrogenase enzyme from Plasmodium falciparum. FEBS J. 274, 2738–2748 (2007).

43. J. Fan et al., Tyrosine Phosphorylation of Lactate Dehydrogenase A Is Important for NADH/NAD+ Redox Homeostasis in Cancer Cells. Mol. Cell. Biol. (2011), doi:10.1128/MCB.06120-11.

44. D. C. Liemburg-Apers, P. H. G. M. Willems, W. J. H. Koopman, S. Grefte, Interactions between mitochondrial reactive oxygen species and cellular glucose metabolism. Arch. Toxicol. (2015), doi:10.1007/s00204-015-1520-y.

45. D. B. Zorov, M. Juhaszova, S. J. Sollott, Mitochondrial Reactive Oxygen Species (ROS) and ROS-Induced ROS Release. Physiol. Rev. (2014), doi:10.1152/physrev.00026.2013.

46. A. A. Nabeebaccus et al., Nox4 reprograms cardiac substrate metabolism via protein O-GlcNAcylation to enhance stress adaptation. JCI insight (2017), doi:10.1172/jci.insight.96184.

47. M. Zhang et al., NADPH oxidase-4 mediates protection against chronic load-induced stress in mouse hearts by enhancing angiogenesis. Proc. Natl. Acad. Sci. (2010), doi:10.1073/pnas.1009700107.

48. L. Viera, Y. Z. Ye, A. G. Estévez, J. S. Beckman, in Methods in Enzymology (1999; http://linkinghub.elsevier.com/retrieve/pii/S0076687999011015), vol. 301, pp. 373–381.

49. A. Görg, W. Weiss, M. J. Dunn, Current two-dimensional electrophoresis technology for proteomics. Proteomics. 4, 3665–3685 (2004).

50. D. H. Perlman, E. A. Berg, P. B. O’Connor, C. E. Costello, J. Hu, Reverse transcription-associated dephosphorylation of hepadnavirus nucleocapsids. Proc. Natl. Acad. Sci. 102, 9020–9025 (2005).

51. M. Karas, F. Hillenkamp, Laser desorption ionization of proteins with molecular masses exceeding 10,000 daltons. Anal. Chem. 60, 2299–2301 (1988).

52. A. W. Sousa da Silva, W. F. Vranken, ACPYPE - AnteChamber PYthon Parser interfacE. BMC Res. Notes. 5, 367 (2012).

53. K. Lindorff-Larsen et al., Proteins Struct. Funct. Bioinforma., in press, doi:10.1002/prot.22711.

54. M. J. Abraham et al., GROMACS: High performance molecular simulations through multi-level parallelism from laptops to supercomputers. SoftwareX. 1–2, 19–25 (2015).

55. W. Humphrey, A. Dalke, K. Schulten, VMD: Visual molecular dynamics. J. Mol. Graph. 14, 33–38 (1996).

56. E. F. Pettersen et al., UCSF Chimera?A visualization system for exploratory research and analysis. J. Comput. Chem. 25, 1605–1612 (2004).

57. P. Pacher, Potent Metalloporphyrin Peroxynitrite Decomposition Catalyst Protects Against the Development of Doxorubicin-Induced Cardiac Dysfunction. Circulation. 107, 896–904 (2003).

